# Diverse environmental perturbations reveal the evolution and context-dependency of genetic effects on gene expression levels

**DOI:** 10.1101/2021.11.04.467311

**Authors:** Amanda J. Lea, Julie Peng, Julien F. Ayroles

## Abstract

There is increasing appreciation that human complex traits are determined by poorly understood interactions between our genomes and daily environments. These “genotype x environment” (GxE) interactions remain difficult to map at the organismal level, but can be uncovered using molecular phenotypes. To do so at large-scale, we profiled transcriptomes across 12 cellular environments using 544 immortalized B cell lines from the 1000 Genomes Project. We mapped the genetic basis of gene expression across environments and revealed a context-dependent genetic architecture: the average heritability of gene expression levels increased in treatment relative to control conditions and, on average, each treatment revealed expression quantitative trait loci (eQTL) at 11% of genes. In total, 22% of all eQTL were context-dependent, and this group was enriched for trait- and disease-associated loci. Further, evolutionary analyses revealed that positive selection has shaped GxE loci involved in responding to immune challenges and hormones, but not man-made chemicals, suggesting there is reduced opportunity for selection to act on responses to molecules recently introduced into human environments. Together, our work highlights the importance of considering an exposure’s evolutionary history when studying and interpreting GxE interactions, and provides new insight into the evolutionary mechanisms that maintain GxE loci in human populations.

## Introduction

It is now clear that most human complex traits and diseases are determined by poorly understood interactions between an individual’s genetic background and his or her environment [1–3]. Consequently, there has been a strong interest in mapping “genotype x environment (GxE) interactions”—in which genotype predicts an individual’s response to environmental variation— and in understanding how loci involved in GxE interactions evolve and are maintained in our species. However, scientists have struggled in practice to map GxE interactions in human populations, largely because 1) the relevant environmental factors are often unknown, difficult to measure, or minimally variable within the study population and 2) large sample sizes are needed to overcome the power limitations posed by point #1. Current state of the art approaches have focused on leveraging very large cohort studies such as the UK Biobank, however, this work has only uncovered a handful of loci involved in GxE interactions for common diseases and complex traits [4–7].

An alternative approach is to use *in vitro* manipulations of the cellular environment paired with transcriptomics to map “context-dependent” expression quantitative trait loci (eQTL), defined as variants that do not affect gene expression levels under baseline conditions but become associated with transcriptional variation following an *in vitro* exposure (or vice versa). This approach focuses on the cellular level as a proxy for the organismal level in a tractable, experimental framework. Using this methodology, thousands of GxE interactions, in the form of context-dependent eQTL, have been identified following cell treatment with pathogens, other molecules that provoke an immune response, drugs, hormones, chemicals, and additional stimuli [8–16]. These studies have consistently shown that context-dependent eQTL overlap genome-wide association (GWAS) hits for complex traits and diseases, in some cases more so that eQTL that are constant or “ubiquitous” across cellular conditions [8,16]. These studies have also revealed that context-dependent eQTL revealed by immune stimuli are often more strongly enriched for signatures of past adaptation than ubiquitous eQTL, suggesting there has been historical selection for plasticity in immune function.

Taken together, this body of work argues that GxE interactions and context-dependent eQTL make important contributions to the genetic architecture of gene expression levels. However, given the large sample sizes needed to robustly map context-dependent eQTL, only a handful of cellular conditions (mostly immune-related) have ever been explored in any single large-scale, genome-wide study. This leaves us with a poor understanding of 1) the full catalog of GxE interactions, as new cellular perturbations typically reveal new context-dependent eQTL [8–16] and 2) the evolutionary forces that maintain context-dependent eQTL. In particular, while previous work has focused on the action of positive selection is maintaining context-dependent eQTL for pathogens and immune stimuli, which have co-evolved with humans through evolutionary time, context-dependent eQTL for “evolutionarily novel” stimuli (e.g., man-made chemicals that have only recently been introduced into our environments) may be maintained by different forces. For example, inefficient purifying selection may maintain GxE loci that are in fact deleterious, but are only revealed under rare or newly introduced environmental conditions; in other words, these loci may be sufficiently “hidden” from purifying selection such that they persist in human populations despite their negative effects. However, empirical investigations of the varied evolutionary forces that maintain context-dependent eQTL, especially in high-powered studies exploring many types of cellular environments, remain extremely limited.

Here, we report a large-scale study in which we profiled genome-wide gene expression levels across 12 different cellular environments using 544 immortalized B cell (lymphoblastoid) lines from the 1000 Genomes Project [17]. We focused on unrelated European and African individuals for whom whole genome sequence data was publicly available [18], and we used these data to map both ubiquitous and context-dependent eQTL across all 12 cellular conditions. Our experimental treatments included stimuli familiar to B cells such as immune signaling molecules and hormones, but also man-made chemicals and other novel cell stressors that have not co-evolved with human B cells through evolutionary time. Our design thus allowed us to address three fundamental questions about the genetic architecture of gene expression levels: 1) to what degree is gene expression determined by ancestry (as has been shown previously [9,12]), and how environmentally robust are these effects? 2) how prevalent are context-dependent versus ubiquitous eQTL, and what are their respective properties and relevance to human complex traits? and (3) what are the evolutionary forces (e.g., genetic drift, inefficient purifying selection, positive selection) that maintain context-dependent versus ubiquitous eQTL, and do these forces differ depending on the evolutionary history of the cell treatment? Together, our work emphasizes the importance of GxE interactions in shaping complex traits, and provides new insight into the evolutionary mechanisms that maintain GxE loci in human populations.

## Results

### Diverse cell exposures induce diverse changes in gene expression levels

We exposed 544 lymphoblastoid cell lines (LCLs) derived from individuals included in the 1000 Genomes study [17] to 12 cellular environments, including 10 treatment and 2 control conditions (Figure 1, Table 1, and Table S1). We chose treatments that have been previously shown to induce moderate to strong responses in LCLs, focusing on a range of treatment types including immune and non-immune-related stimuli. Specifically, we exposed cells to: 1-2) FSL-1 and gardiquimod, two synthetic molecules that activate the TLR2/TLR6 and TLR7 signaling pathways, respectively; 3) B-cell-activating factor, a cytokine that is a member of the tumor necrosis factor family and a potent B cell activator; 4) interferon gamma, a cytokine that is critical for coordinating innate and adaptive immune responses to viral infections; 5) dexamethasone, a synthetic glucocorticoid hormone and anti-inflammatory drug; 6) insulin-like growth factor 1, a hormone that plays a key role in growth-related processes; 7) tunicamycin, an antibiotic that induces endoplasmic reticulum stress and is used as a model for cell stressors that impact protein folding; 8-10) perfluorooctanoic acid, acrylamide, and bisphenol A, three man-made chemicals and environmental contaminants; 11) ethanol, a vehicle control for tunicamycin and dexamethasone, but also considered a treatment with water as a control; and 12) water, a vehicle control for all treatments except tunicamycin and dexamethasone; see Table S2). Following 4 hours of exposure to each cellular environment, we extracted RNA and used TM3’seq [19] to collect mRNA-seq data from 5223 samples (mean reads per sample ± SD = 2.199 ± 2.731 million). We paired this mRNA-seq data with publicly available genotype data for the same individuals, derived from whole genome sequencing to at least 30x coverage [18]. All individuals included in our study were unrelated and exhibited ancestry of European or African origin (admixed populations were not included in our study design; Figure 1). After filtering, we retained a total of 3886 mRNA-seq profiles from 500 unique individuals (Table S3). Genotype data derived from high coverage whole genome sequencing was also available for 454 of these individuals (Table S4).

**Figure 1.**
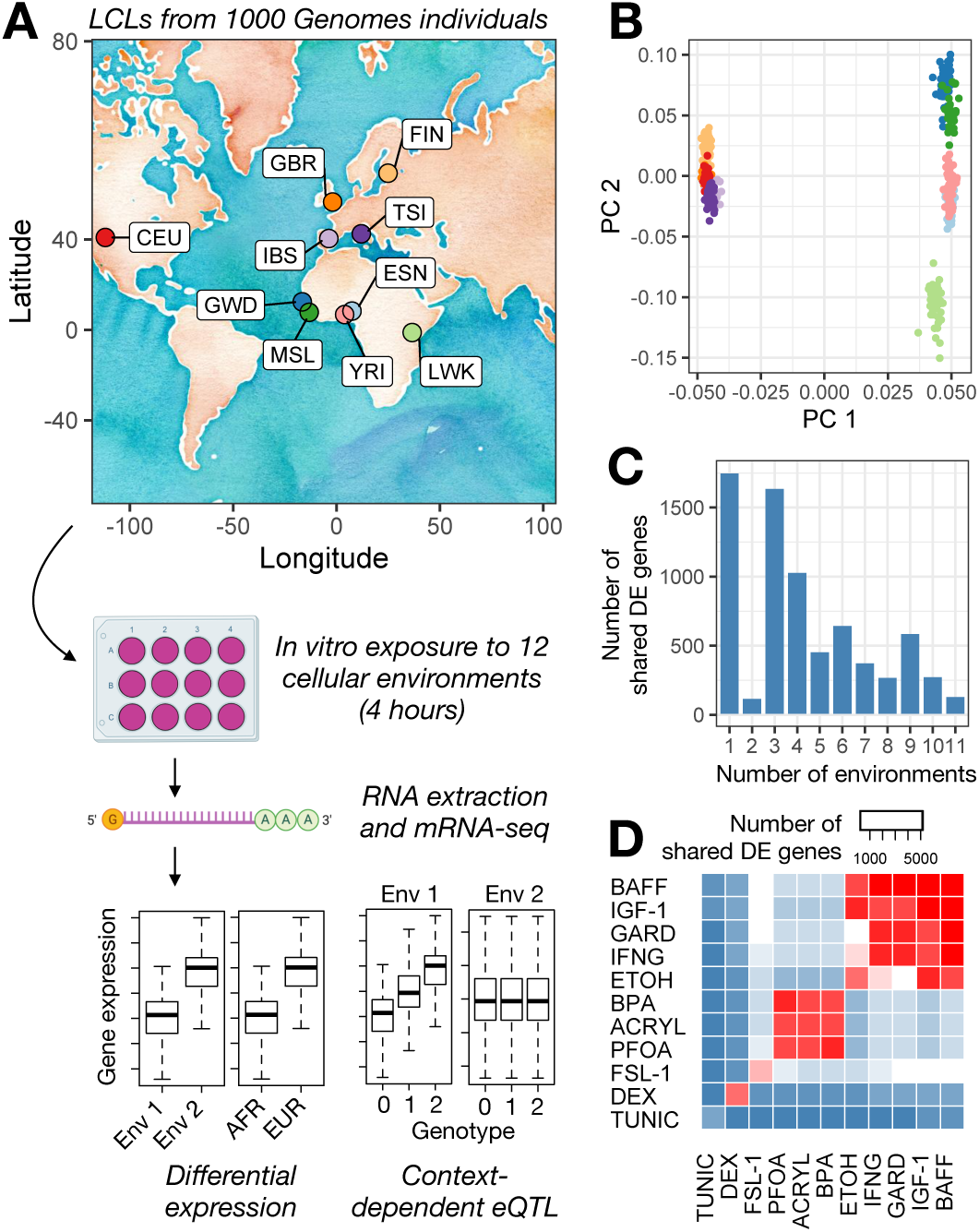
Study overview and environmental effects on gene expression. (A) Lymphoblastoid cell lines (LCLs) derived from individuals included in the 1000 Genomes Project were obtained from Coriell Institute. Specifically, we obtained the lines listed in Table S1, which were derived from individuals of European and African ancestry as noted on the map (abbreviations for included populations are as follows: CEU=Utah residents (CEPH), GWD=Gambian Mandinka, GBR=British, IBS=Iberian, MSL=Mende, FIN=Finnish, TSI=Toscani, ESN=Esan, YRI=Yoruba, LWK=Luhya). Each cell line was exposed to 12 cellular environments for 4 hours, after which we harvested the RNA and performed mRNA-seq. These data were used to understand differential expression as a function of environmental context and ancestry (AFR=African, EUR=European), as well as to map ubiquitous and context-dependent eQTL. (B) Principal components analysis of genotype data for individuals included in this study (colors are as in A). Individual used in this analysis are those for which paired RNA-seq and genotype data were available (Table S4). (C) Number of differentially expressed (DE) genes shared between N environments using a mashR, joint analysis approach. N is plotted on the x-axis and ranges from 1 (i.e., the gene is DE in response to only 1 environmental treatment) to 11 (i.e., the gene is DE in response to all 11 environmental treatments). The low number of genes when N=2 is driven by a large number of DEX-specific genes, such that 93.7% of the N=1 genes are DEX-specific; when DEX is excluded from the dataset, most genes are shared between three or more environments (Figure S5). (D) Number of differentially expressed (DE) genes shared between a given pair of environmental treatments using a mashR, joint analysis approach. The diagonal represents the number of DE genes in response to the focal environmental treatment.

**Table 1.** Cellular environments.

| Environment | Abbreviation | Treatment category |
| --- | --- | --- |
| Gardiquimod | GARD | Immune stimulant |
| Fibroblast-stimulating lipopeptide 1 | FSL-1 |  |
| Interferon gamma | IFNγ | |
| B cell activating factor | BAFF |  |
| Dexamethasone | DEX | Hormone |
| Insulin-like growth factor 1 | IGF-1 |  |
| Acrylamide | ACRYL | Novel contaminant or cell stressor |
| Perfluorooctanoic acid | PFOA |  |
| Bisphenol A | BPA |  |
| Tunicamycin | TUNIC |  |
| Ethanol | ETOH |  |
| Water | H2O | Vehicle control |

We found significant differences in transcriptome dynamics between treated and control cells (when analyzing each of the 11 treatment-control pairs separately): on average, 6.39% ± 8.98% (SD) of all 10157 tested genes responded to a given treatment, with a maximum of 31.64% genes differentially expressed in response to dexamethasone (limma FDR<10%; Figure 1 and Table S5). Gene set enrichment analyses (GSEA) revealed significant overrepresentation of differentially expressed genes in expected biological pathways. For example, gardiquimod induced differential expression of genes involved in immune system processes such as “cytokine production” (enrichment score=0.536, q=0.055), “toll like receptor signaling pathway” (enrichment score=0.507, q=1.03×10^−2^), and “activation of immune response (enrichment score=0.457, q=1.03×10^−2^). Similarly, IFNγ treatment activated expression of genes related to “response to virus” (enrichment score=0.583, q=0.066), “type I interferon production” (enrichment score=0.540, q=0.066), and “regulation of innate immune response” (enrichment score=0.462, q=0.066). Finally, the strongest pathways induced by tunicamycin were related to endoplasmic reticulum stress, such as “endoplasmic reticulum unfolded protein response” (enrichment score=0.581, q=0.189) and “ER overload response” (enrichment score=0.844, q=0.189; see Figure S1 and Table S6 for results for all treatments). These results suggest that our experimental cell treatments induced appropriate biological responses.

When we used an empirical Bayes approach to perform joint analyses across our entire dataset [20], we found that 23.93% of differentially expressed genes were unique to a single treatment (mashR LFSR<10% and no posterior effect size estimates within a factor of 2). In contrast, 42.44% were shared between 2 and 10 treatments and a small subset of genes—1.85% of genes differentially expressed in response to any treatment—were generally environmentally responsive and differentially expressed in response to all 11 treatments (mashR LFSR<10% and all posterior effect size estimates within a factor of 2; see Figure 1, Figure S2, and Table S7). These results emphasize the environmental dependency of gene expression, as well as the fact that diverse experimental treatments provoke non-overlapping cellular responses.

### Genetic ancestry controls the expression of immune-related genes across cellular conditions

We next explored transcriptional variation as a function of ancestry, comparing individuals of African versus European descent (Figure 1). Of the 10157 genes we tested, an average of 3.59% ± 2.80% (SD) genes were differentially expressed between ancestry groups (limma FDR<10%; Table S8). Joint analyses [20] revealed that these ancestry-associated genes were generally shared across conditions, with 69.20% of ancestry-associated genes displaying similar effect sizes across all 12 conditions, and 98.92% displaying similar effect sizes across ≥2/3 conditions (Figure 2, Figure S2, and Table S7). In agreement with previous work [9], we found that genes controlled by ancestry were most strongly enriched for immune system processes such as “cytokine-mediated signaling pathway” (enrichment score=0.311, q=0.160), “response to virus” (enrichment score=0.419, q=0.160), and “inflammatory response” (enrichment score=0.350, q=0.160; Figure 2 and Table S9). Further, we downloaded publicly available sets of genes that are thought to lie along the causal pathway linking genetic variation, gene expression, and human complex traits (inferred through Probabilistic Transcriptome Wide Association Studies, PTWAS [21]) and tested for overlap with our set of ancestry-associated genes. Here, we found that ancestry-associated genes were enriched for complex traits and diseases with immune involvement, such as lymphocyte counts (fold enrichment=1.10, q=0.098), platelet counts (fold enrichment=1.08, q=0.098), inflammatory bowel disease (fold enrichment=5.59, q=0), and Crohn’s disease (fold enrichment =5.59, q=0) (Table S10). Importantly, many of these phenotypes are known to differ in prevalence and/or etiology between individuals of African versus European ancestry [22,23].

**Figure 2.**
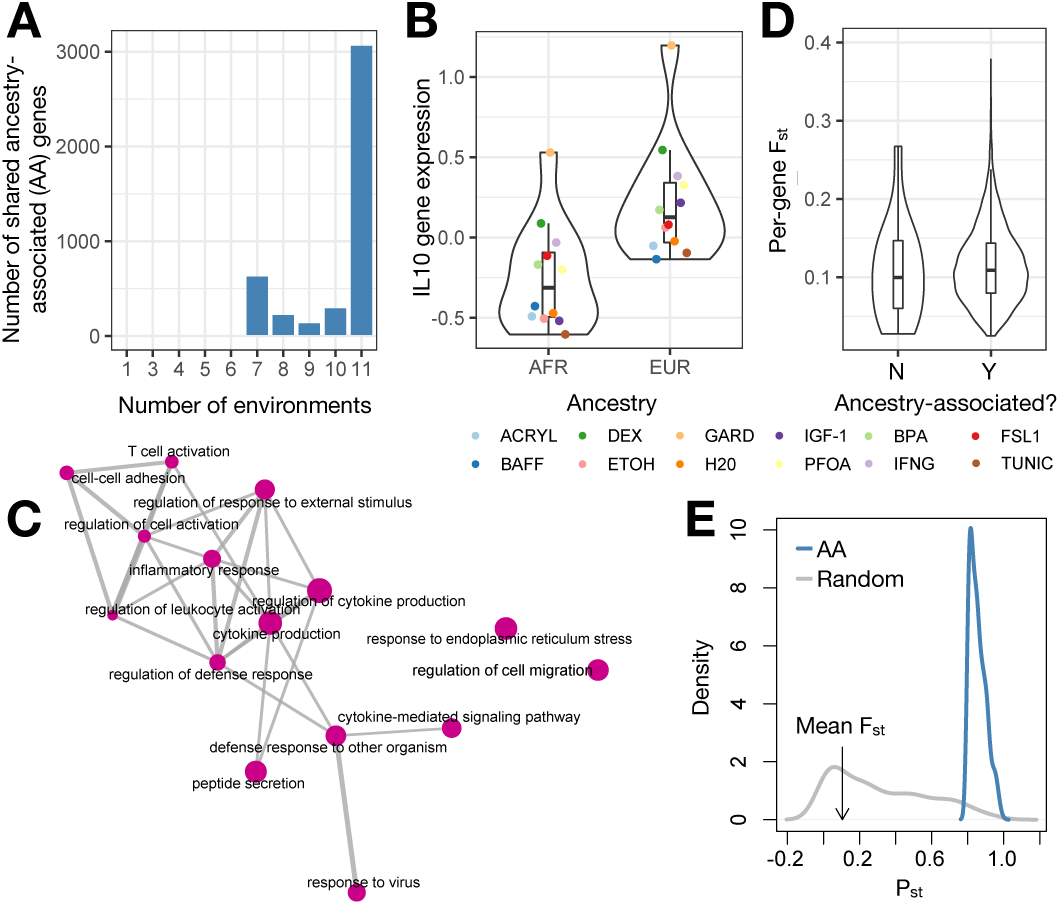
Ancestry effects on gene expression. (A) Number of ancestry-associated (AA) genes shared between N environments using a mashR, joint analysis approach. N is plotted on the x-axis and ranges from 1 (i.e., the gene is AA in only 1 cellular environment) to 12 (i.e., the gene is AA in all 12 cellular environments). (B) Example of an ancestry-associated gene. Y-axis shows the mean, normalized *IL10* gene expression levels estimated in each environment, after regressing out 3 surrogate variables. (C) Results from gene set enrichment analyses testing for overrepresentation of particular gene ontology categories among AA genes (note, genes were sorted by average AA effect size across all 12 cellular environments and only the top 15 most significant categories are shown). Enrichment map was created with the emapplot function in the R package enrichplot. (D) Distribution of average per-gene F_ST_ values for genes that 1) were found to be AA in most (>2/3) cellular environments or 2) had no effects on ancestry in any cellular environment (results are from a mashR, joint analysis approach). (E) Phenotypic differentiation (in gene expression; P_ST_) versus genetic differentiation (F_ST_) for AFR versus EUR samples. Plots show the distribution of P_ST_ values for: 1) AA genes identified in the H20 cellular environment (blue) and 2) a same-sized set of randomly selected genes (grey). The mean genome-wide F_ST_ value comparing genetic divergence between AFR and EUR samples is noted on the x-axis with an arrow. We find that all AA genes exhibit P_ST_ > F_ST_, indicative of diversifying selection [24,25].

We hypothesized that the observed transcriptional differentiation between African and European individuals was controlled by genetic variation [9,12], and performed two sets of analyses to address this possibility. First, we followed the approach of [9] and asked whether genes with ancestry effects exhibited higher FST values between African and European populations relative to non-ancestry-associated genes. We found that this was indeed the case for genes with shared effects across ≥2/3 of conditions (Wilcoxon signed-rank test, p=1.91×10^−4^) as well as for the total set of genes with effects in any condition (Wilcoxon signed-rank test, p=2.79×10^−4^; Figure 2). Further, when analyzing genes with ancestry effects in ≥1 condition, we found that the degree of genetic differentiation was positively correlated with the number of conditions in which an ancestry effect was found (linear model, beta=3.69×10^−4^, p=7.85×10^−5^), suggesting that genes with ancestry effects that are unmodified by treatment may be under the strongest genetic control. Second, we compared PST values for ancestry-associated genes identified in ≥1 condition versus genome-wide FST values to understand whether ancestry-related variation in the transcriptome (that is putatively genetically controlled) showed evidence for being driven by genetic drift (PST = FST), diversifying selection (PST > FST), or stabilizing selection (PST < FST) [24,25]. Our analyses point toward diversifying selection at ancestry-associated genes (all PST > FST), suggesting that there has been selection for different local optima in African versus European populations (Figure 2 and Figure S3).

Consistent with the high degree of effect size similarity we observed for ancestry effects analyzed across cellular environments, we did not find any compelling evidence for ancestry x treatment effects on gene expression levels using several analysis approaches (see Methods). While previous work has found that ancestry predicts the magnitude of the response to experimental infection in macrophages and peripheral blood mononuclear cells (PBMCs) [9,12,26], we note that our treatments induced much smaller overall transcriptional shifts in LCLs relative to these previous studies in primary cells, which likely affects our power to detect interaction effects at genome-scale. Further, LCLs are derived from B cells which are a key component of the adaptive immune system; in contrast, previous work has found that ancestry effects on the response to infection largely involve innate immune system cell types and processes [9,12,26].

### Cellular perturbations reveal many context-dependent eQTL

The primary goal of this study was to understand the genetic architecture of gene expression levels, including the degree to which genetic effects are context-dependent versus unperturbed by environmental challenges. Consistent with previous studies [27–29], we found that there is a substantial genetic component to gene expression levels in lymphoblastoid cell lines, with an average heritability of 0.111 ± 0.250 (SD) in unstimulated cells. However, we found that this genetic component changed following cell treatments, such that mean per-gene heritability estimates were significantly higher in almost all treatment conditions relative to their respective controls. Specifically, mean heritability estimates increased in 9 of 11 treatments, with an average fold increase of 1.736 ± 0.955 (SD) across all 11 treatments (Figure 3 and Table S5). Further, the difference in mean per-gene heritability estimates between treatment and control conditions remained after accounting for the sample size used to estimate heritability in each condition (linear model: β=2.94×10^−2^, p=2.90×10^−13^) and after subsampling all environments to an identical sample size (Figure 3 and Table S5). Together, these results are consistent with an increase in additive genetic variation for gene expression levels when cells are perturbed.

**Figure 3.**
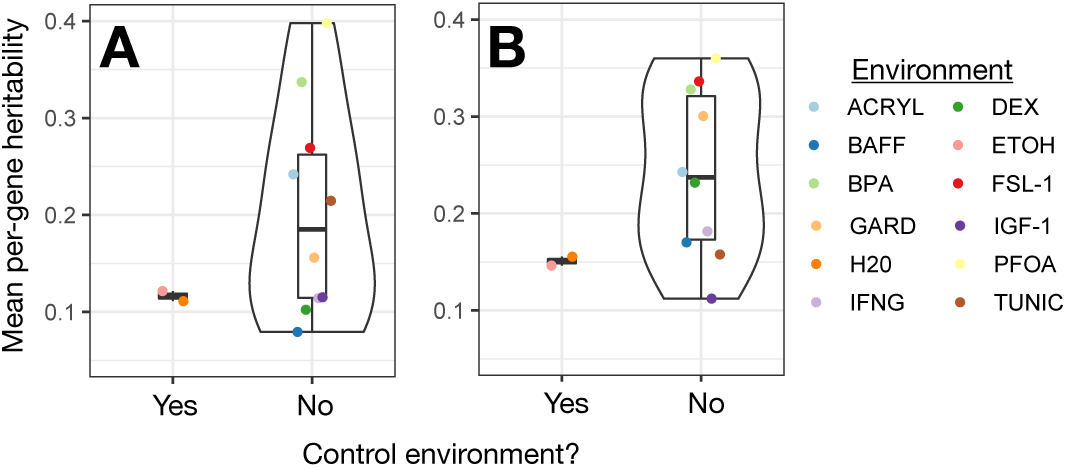
Environmental perturbations increase the heritability of gene expression levels. Y-axis shows the mean per-gene heritability estimated in each environment, using (A) the total available sample size for each environment or (B) a subsample of n=100 from each environment (plot shows the average of 5 subsamples).

We confirmed that the genetic architecture of gene expression levels changes following cellular perturbations by mapping cis eQTL in each of the 12 control or treatments conditions and comparing their effect sizes. These analyses revealed extensive genetic control of gene expression: 11.07% ± 7.64% (SD) of genes contained at least one eQTL (matrixeQTL FDR<10%), with this number increasing to 15.72% ± 3.69% (SD) when only considering the conditions with the largest sample sizes (Table S8). Further, when we pooled our data and performed one analysis using samples from all conditions, we found that almost all genes (92% of those tested) had at least one eQTL (matrixeQTL FDR<10%).

When we performed joint analyses [20] to identify ubiquitous and context-dependent eQTL, we found that 77.53% of significant eQTL were shared across all 12 conditions, while 6.70% were condition-specific and the remaining 15.78% were shared between 2 and 11 conditions (mean=10.40 ± 3.42 (SD) conditions; Figure 4 and Table S7). We also found substantial evidence for a subset of context-dependent eQTL known as “response eQTL”, which we define as SNPs that do not affect gene expression in the control condition but for which genetic effects are revealed by experimental treatment (or vice versa; Table S5). On average, we found that 11.28% ± 2.91% (SD) of all tested genes contained at least one response eQTL for a given treatment-control pair, with the strongest evidence for response eQTL observed for the treatment that induced the largest overall shift in transcriptome dynamics (i.e., dexamethasone, which revealed response eQTL at 14.98% of genes).

**Figure 4.**
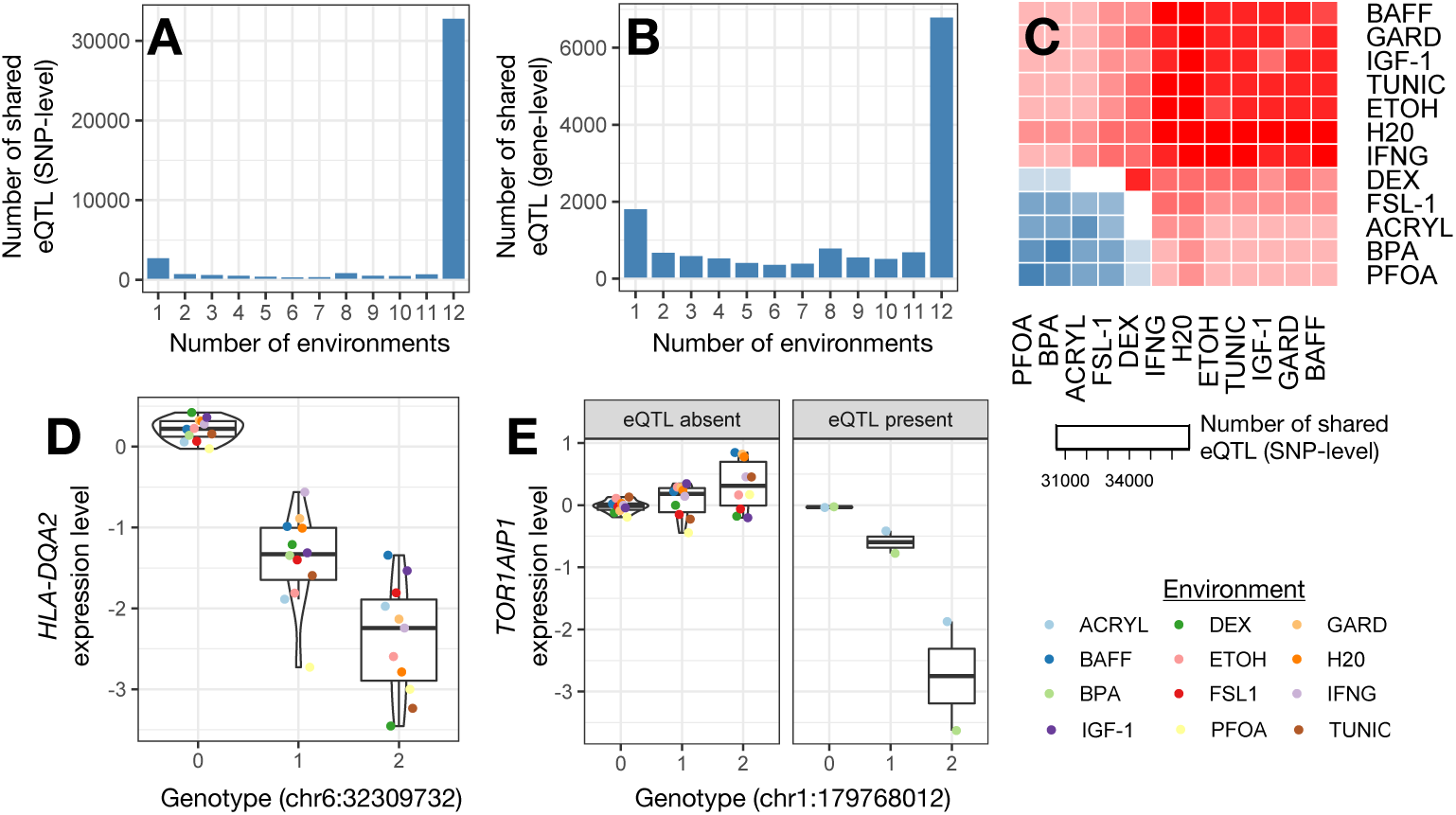
Environmental variation reveals context-dependent eQTL. (A) Number of eQTL shared between N environments using a mashR, joint analysis approach. N is plotted on the x-axis and ranges from 1 (i.e., the eQTL is present in only 1 cellular environment) to 12 (i.e., the eQTL is present in all 12 cellular environments). (B) Same plot as in A, but sharing is defined at the gene rather than the SNP level. (C) Number of eQTL shared between a given pair of cellular environments using a mashR, joint analysis approach. The diagonal represents the number of eQTL in the focal cellular environment. Note that ACRYL, BPA, PFOA, and FSL-1 had lower sample sizes than the other 8 environments, and their clustering thus reflects differences in power. (D-E) Examples of ubiquitous and context-dependent eQTL, identified using a mashR, joint analysis approach. Y-axis shows the mean, normalized expression levels for a given gene estimated in each environment, after regressing out 3 surrogate variables.

To validate and contextualize the eQTL we identified, we performed two follow up analyses: 1) we tested for an expected mechanistic pattern, namely enrichment of eQTL within accessible chromatin regions and 2) we tested for overlap between our eQTL containing genes (“eGenes”) and eGenes identified by The Genotype-Tissue Expression (GTEx) Project in untreated LCLs [30]. First, as others have observed [9,30], we found that SNPs within accessible chromatin regions were more likely to be identified as ubiquitous (p=0.042, hypergeometric test) and context-dependent eQTL (p=4.01×10^−4^; enrichment analyses performed using publicly available ATAC-seq data [31] from untreated LCLs [32]). Second, we found that genes containing ubiquitous eQTL overlapped significantly with GTEx LCL eGenes (p=6.22×10^−3^, hypergeometric test), suggesting there is a core set of eQTL identified across study designs and cell states. However, genes containing context-dependent eQTL did not overlap with GTEx LCL eGenes (p=0.465), suggesting that our cellular perturbations revealed previously uncharacterized genetic effects on gene expression. In support of this idea, genes with evidence for eQTL in our dataset, but not in GTEx, were enriched for genes that responded to at least one experimental treatment (p=4.66×10^−4^).

### Phenotypic relevance of ubiquitous and context-dependent eQTL

Our next goal was to understand the phenotypic relevance of context-dependent versus ubiquitous eQTL, and to test the hypothesis that context-dependent eQTL are especially important for human diseases and adaptively relevant traits. To do so, we asked whether our context-dependent and ubiquitous eQTL SNP sets (or their associated genes) were enriched within three publicly available datasets: 1) GWAS hits for human traits and diseases [33], 2) sets of genes that are thought to lie along the causal pathway linking genetic variation, gene expression, and 114 complex traits and diseases (inferred through PTWAS [21]) and 3) loss of function, mutation-intolerant genes (as curated by ExAC [34]).

First, we found that both context-dependent and ubiquitous eQTL were strongly enriched for loci previously implicated in GWAS of human complex traits and diseases, though contrary to our expectations, these results were slightly stronger for ubiquitous compared to context-dependent eQTL (p-value=5.74×10^−15^ and <10-^16^ for context-dependent and ubiquitous eQTL, respectively, hypergeometric test; Figure 5). Second, we found that context-dependent eGenes were enriched for 11 complex traits, largely focused on immune system traits and disorders such as platelet counts (q=2.47×10^−4^, hypergeometric test), neutrophil counts (q=0.086), and rheumatoid arthritis (q=0.086; Table S10). In contrast, ubiquitous eGenes were enriched for approximately half the number of complex traits (n=6), again including many immune phenotypes such as inflammatory bowel disease (q<10^−16^) and lymphocyte counts (q=0.053; Figure 5 and Table S10).

**Figure 5.**
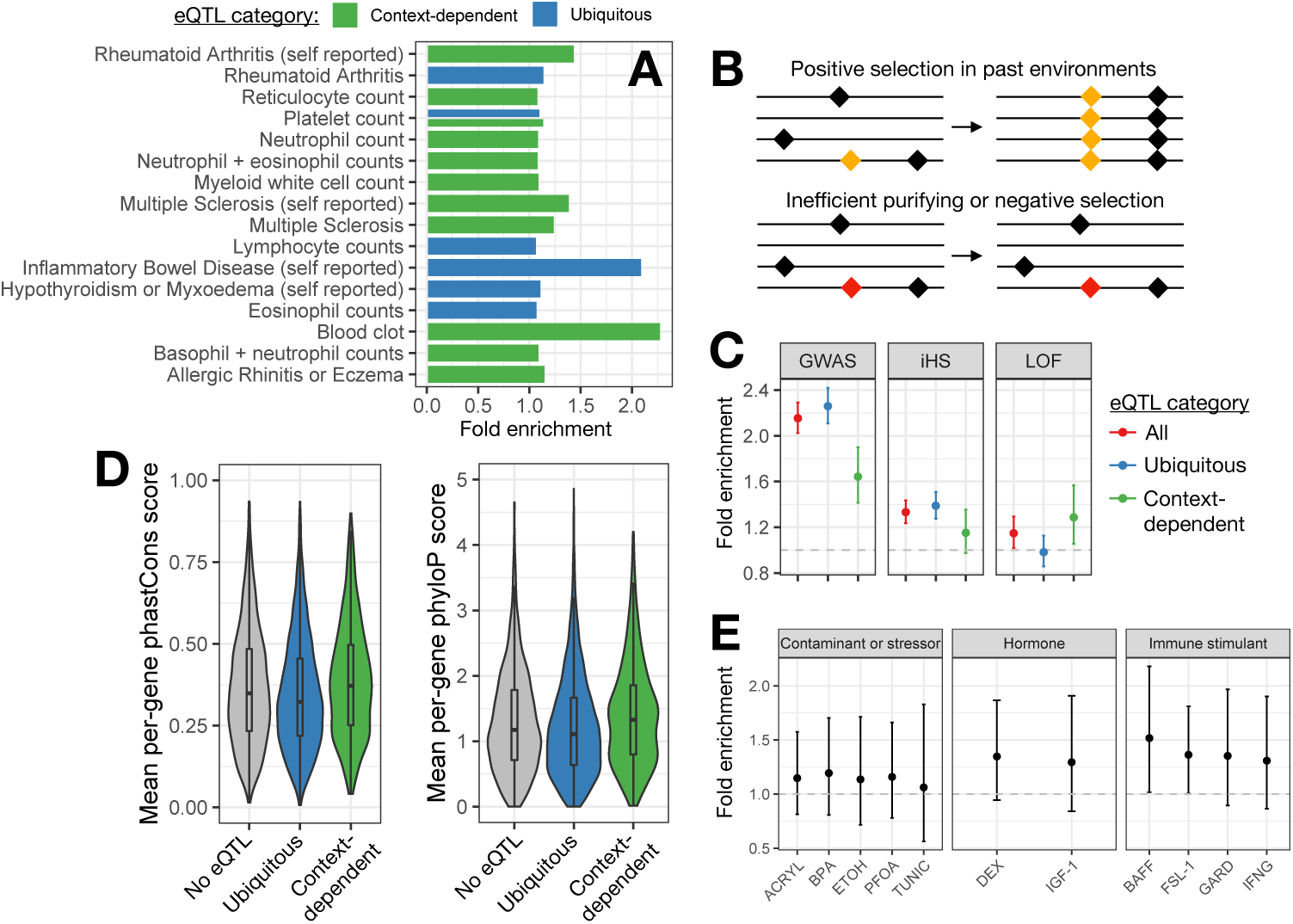
Phenotypic relevance and evolution of context-dependent eQTL. (A) Diseases and heath conditions for which associated genes (identified via PTWAS [21]) are enriched among genes with ≥1 context-dependent or ubiquitous eQTL. X-axis represents the fold enrichment estimate from a Fisher’s exact test. (B) Evolutionary forces that potentially maintain context-dependent eQTL: 1) positive selection on beneficial mutations (yellow) or 2) inefficient purifying or negative selection that fails to remove deleterious mutations (red). (C) Overlap of ubiquitous, context-dependent, and all (ubiquitous and context-dependent) eQTL with 1) the full catalog of GWAS-associated loci [33], 2) iHS outlier loci [36], and 3) genes annotated as loss of function, mutation intolerant [34]. Y-axis represents fold enrichment from a Fisher’s exact test, with error bars denoting the 95% confidence interval for each estimate. (D) Distribution of mean per-gene phastCons and phyloP scores for genes with no eQTL, ≥1 ubiquitous eQTL, and ≥1 context-dependent eQTL. (E) Overlap of response eQTL, identified in a given condition, with iHS outlier loci [36]. Y-axis represents fold enrichment from a Fisher’s exact test, with error bars denoting 95% confidence intervals.

Finally, we tested whether ubiquitous or context-dependent eGenes were enriched for mutation-intolerant genes, which are thought to be phenotypically relevant and essential for life given that no adult humans (in a large population sample known as ExAC) contain mutations that would result in a truncated or loss of function protein [34]. Here, we found that context-specific eGenes were enriched among mutation-intolerant genes (odds=1.306, p=5.45×10-8, Fisher’s exact test), but ubiquitous eGenes were not, and even trended toward being under enriched (odds=0.982, p=0.8102; Figure 5). These results parallel those obtained by the GTEx project: eGenes that were shared across many tissues were significantly depleted for loss of function, mutation-intolerant genes, while tissue-specific eGenes were significantly enriched [30]. These results were interpreted as evidence for purifying selection on regulatory variants that involve many tissues and are thus more likely to have deleterious, pleiotropic effects. In contrast, tissue-specific eQTL, and in our case, context-dependent eQTL, appear to be more likely to “escape” the effects of purifying selection and to persist in loss of function, mutation-intolerant genes.

### Evolutionary forces maintaining ubiquitous and context-dependent eQTL

The analyses described in the previous section point toward negative selection as a key evolutionary force that patterns genetic effects on gene expression. However, several previous studies of context-dependent eQTL uncovered by immune stimulation have instead emphasized a role for positive selection [9,12,16,26]. For example, Kim-Helmuth and colleagues found that monocyte eQTL with differential effect sizes in baseline versus immune-stimulated conditions were enriched among genomic regions with recent signatures of positive selection; a similar result was found for ubiquitous eQTL, though the enrichment was weaker [16]. Drawing on this work and our results thus far, we were motivated to further explore the role of both positive and negative selection in generating context-dependent and ubiquitous eQTL.

To do so, we first drew on publicly available measures of sequence conservation across mammals, to understand whether context-dependent and ubiquitous eQTL fall in rapidly evolving versus conserved regions of the human genome. We found that ubiquitous eGenes exhibited both lower phastCons (p=1.765×10^−5^, Wilcoxon signed-rank test) and phyloP scores (p=1.749×10^−4^) than background expectations, indicating that ubiquitous eGenes are underrepresented in evolutionarily conserved genes. In contrast, context-dependent eGenes exhibited higher phastCons (p=5.01×10^−3^, Wilcoxon signed-rank test) and phyloP scores (p=1.35×10^−4^; Figure 5) than background expectations, indicating that they are overrepresented in conserved genes. Similar to our results using mutation-intolerant gene annotations, these analyses suggest that, in highly conserved and putatively essential genes, ubiquitous eQTL are selected against and depleted, while context-dependent eQTL are “hidden” from selection and persist.

Next, we explicitly investigated a role for positive selection by obtaining per-site estimates of the integrated haplotype score (iHS), a commonly used measure of within-population recent positive selection [35,36]. We obtained genome-wide iHS estimates for each of the populations included in our study, and identified putative selection candidates as loci that fell in the >99^th^ percentile of |iHS| values in ≥2 populations (as in [9,16]). When we overlapped these selection outliers with our eQTL, we found that loci that functioned as eQTL (in all or <12 cellular environments) were more likely to show signatures of positive selection (odds=1.332, p=3.702×10^−13^, Fisher’s exact test). This result was especially strong for ubiquitous eQTL (odds=1.389, p=1.688×10^−13^), but less so for context-dependent eQTL (odds=1.152, p=0.086; Figure 5), in contrast to patterns observed in previous work focusing on immune stimulations [9,16]. Given that our study focuses on a more diverse set of environmental perturbations, including immune stimulations but also hormones, environmental contaminants, and evolutionarily novel cell stressors, we wondered whether heterogeneity between treatments could explain the lack of enrichment of iHS outliers in the total set of context-dependent eQTL. When we analyzed each condition separately, we found significant enrichment of iHS outliers in response eQTL for several immune stimuli, namely FSL-1 (odds=1.362, p=0.036) and BAFF (odds=1.512, p=0.031). Further, we found that enrichment effect sizes were consistently higher for the immune treatments relative to the environmental contaminant and cell stressor treatments (β=-0.246, p=5.82×10^−4^, linear model) but not the hormone treatments (β=-0.065, p=0.297; Figure 5). Thus, our data suggest that positive selection has played a much greater role in shaping loci involved in the response to immune signaling molecules and hormones relative to loci involved in the response to man-made chemicals and novel cell stressors.

## Discussion

Despite major advances in genomic technologies, understanding the genetic basis of human complex traits remains a challenge. While decades of GWAS have uncovered many loci for common diseases and health-related traits, there is a general consensus that mapping additive effects will not allow us to account for the total estimated genetic component of most phenotypes. This problem is known as the “missing heritability” problem, and has been used to argue for a critical role for GxE interactions [37]. More specific support that GxE interactions contribute to human complex traits comes from analyses of variance and observations that the heritability of key traits has increased in recent decades, despite minimal changes in the genetic makeup of populations [38–41]. However, despite multiple lines of evidence that GxE interactions are likely important, we have made little progress in mapping the specific loci involved in GxE interactions. This reality limits our ability to understand their distribution, effect sizes, mechanisms of action, and evolution.

Here, we use a cell culture model to overcome issues faced by observational studies and to maximize our power to detect GxE interactions in the form of context-dependent eQTL. Specifically, we exposed cells from genetically well-characterized individuals to 12 controlled, *in vitro* environments and asked whether genetic variation predicted individual responses at the transcriptional level. We found that diverse environmental perturbations induced diverse gene regulatory programs (Figure 1), while the genetic control of gene expression levels was more environmentally robust: at least 70% of ancestry effects on the transcriptome were consistent in effect size across environments, as were 78% of eQTL. These results agree with previous work using smaller sample sizes, fewer environments, as well as cross-tissue comparisons [9,12,15,30].

Nevertheless, our experiments do show that a non-negligible portion of the transcriptome’s genetic architecture is environmentally sensitive: 16% of eQTL were only shared between 2 and 11 conditions, while 6% were specific to a single condition. Interestingly, we found that our observed patterns of cis eQTL sharing generally mimicked the bimodal distribution observed by GTEx for cross-tissue comparisons [30]. In other words, the largest categories of cis eQTL were those shared across all 12 environments, followed by those that were specific to a single environment (Figure 4). Importantly, many of the eQTL revealed by our environmental treatments were both uncharacterized (e.g., unannotated in GTEx LCLs [30]) and putatively phenotypically relevant (Figure 5), arguing that environmental diversity should be incorporated into eQTL mapping studies whenever possible (e.g., [15]). In fact, while an estimated 88% of all genetic variants associated with complex human traits and diseases lie outside of protein-coding regions [42,43], the most comprehensive eQTL studies to date (e.g., GTEx [30]) have only accounted for ∼1/2 of known regulatory GWAS hits. This state of affairs has motivated recent calls to expand the set of cellular conditions under which we study links between genotype, gene regulation, and disease [44]; our study design provides one feasible avenue for doing so.

While previous work has applied *in vitro* environmental manipulations to study GxE interactions [8–16], the number of environments we assay here using genome-wide datasets and appreciable sample sizes is unprecedented. Further, previous experiments have focused heavily on environmental manipulations involving pathogens and other immune stimuli, and we speculate that context-specific eQTL in the immune system may manifest and evolve in different ways than context-specific eQTL for other stimuli, especially stimuli that have not been common throughout human evolutionary history. According to evolutionary theory, most new mutations are overall deleterious (e.g., because of their pleiotropic effects) and will be selected against; in contrast, context-dependent regulatory mutations can provide more targeted fitness benefits, and are thus thought to play a key role in the evolution of adaptively relevant trait [44– 46]. In support of this argument, several studies have shown that eQTL that control the gene regulatory response to infection are under positive selection [9,12,16,26]. However, it is unlikely that natural selection has similarly shaped eQTL that control the response to stimuli recently introduced into human environments (e.g., man-made chemicals). In line with this thinking, we find essentially no evidence that positive selection has shaped GxE loci revealed by novel cell stressors or chemicals, but we do find evidence for this pattern for immune-responsive loci (Figure 5). Our results thus provide novel insight into the fact that cellular perturbations with distinct evolutionary histories can produce divergent patterns. Moving forward, we argue for greater consideration of the evolutionary history of a given environmental exposure when studying and interpreting GxE interactions.

There are several limitations to the present study, as well as open directions for future work. First, our sample sizes were more than reasonable for eQTL mapping in all 12 cellular environments, however, they were not identical. Nevertheless, our use of the mashR framework should circumvent many of these issues via joint analysis and consideration of effect sizes rather than significance cutoffs [20]. Second, our study design relied on Epstein-Barr virus transformed lymphoblastoid cell lines, because they are a replenishable and shareable resource and are commercially available for genetically well-characterized individuals [17]. They have also been used extensively for functional genomic work [47–51], and previous studies have shown that 1) genomic results from LCLs replicate in primary tissues [47–51] and 2) gene expression levels in newly established LCLs maintain a strong individual signature [52]. However, gene regulation in LCLs is not identical to their progenitor B cells, and the transformation process is known to induce certain artifacts [53,54]. Thus, while we believe LCLs represent a powerful model for high-throughput and large-scale GxE mapping, future work in primary tissues or induced pluripotent stem cells will be important for corroborating the patterns we see here. Finally, we did not generate environment-specific data on additional gene regulatory mechanisms such as DNA methylation, chromatin accessibility, or chromatin state. While this work was beyond the scope of the present study, it is a key avenue for future research and for understanding the molecular mechanisms that generate context-dependent eQTL (e.g., environmentally dependent transcription factor binding). While functional fine-mapping of individual GxE interactions can be laborious and difficult, several recent studies offer promising new avenues for using gene regulatory assays to understand the path from genotype to phenotype [55,56], including under diverse environmental conditions.

Technological advances have fueled the ascent of personal genomics and the promise of precision medicine. However, to unlock this potential, we must first understand how the environmental and genetic interactions unique to each individual contribute to variation in complex traits. Our study provides a comprehensive window into the environmental-dependency of the human transcriptome, and highlights that diverse environmental exposures leads to an array of unique cellular responses. Importantly, these responses are modulated by an individual’s genetic background, including by genetic effects that are only revealed by environmental change (a class of variants known as “cryptic genetic variation” [57,58]). Cryptic alleles likely drive the significant increase in heritability we observe following cellular treatments, and could play a major role in explaining the “missing heritability” problem; again, these results argue for a critical role for GxE interactions in driving variation in complex traits.

## Materials and Methods

### Cell culture, experimental cell treatments, and mRNA-seq

Lymphoblastoid cell lines (LCLs) were obtained for 544 unrelated individuals included in the 1000 Genomes study [17]. All cells were ordered from Coriell Institute, and live cultures were shipped overnight to Princeton University in randomized batches of 25 (Table S1). Cells were cultured in parallel for 5-11 days until 12 million cells were available to seed (at a density of 1 million cells/2.5mL of media in a 12 well plate). After an overnight incubation period, 12 environmental treatments (Table 1 and Table S2) were added to each of the 12 wells (see for treatment concentrations). After 4 hours, cells were washed, harvested, and preserved in lysis buffer for downstream RNA work.

Total RNA was extracted from each sample using Zymo’s Quick-RNA 96 kit, following the manufacturer’s instructions. mRNA-seq libraries were prepared using the published TM3’seq protocol [19] and a CyBio FeliX liquid handling robot (Analitik Jena). The total dataset (n=5223 libraries) was sequenced across four runs of the Illumina NovaSeq platform. Each sample was sequenced to a mean depth of 2.199 ± 2.731 (SD) million reads using 100bp single end sequencing. Additional information relating to cell culture, experimental cell treatments, and mRNA-seq data generation is available in the Supplementary Materials.

### Low level processing of mRNA-seq and genotype data

Following sequencing, we trimmed each FASTQ file for low quality bases and adapter contamination [59], mapped the filtered reads to the human reference genome (hg38) [60], and extracted counts of reads mapped to genes [61]. If a sample had fewer than 250,000 reads mapped to protein coding genes, we excluded it from further analyses (see list of filtered samples in Table S3). We further filtered the per-gene counts matrix to exclude lowly expressed genes, normalized the count data [62], and conducted surrogate variable (SV) analysis to remove variance attributed to batch or technical effects [63] (specifically, we fit three SVs and then regressed out their effects). Additional information relating to low level processing of mRNA-seq data is available in the Supplementary Materials.

We downloaded phased genotype calls, derived from ∼30x whole genome sequence data, for 454 1000 Genomes Project individuals included in our study [18] (Table S4). We the used Plink [64] to remove the following variant types: indels, SNPs with >2 alleles, SNPs with MAF<0.05, SNPs called in <50% of individuals, and SNPs out of Hardy-Weinberg equilibrium (p<10^−6^). This filtering left us with 7,205,828 SNPs. We then performed LD filtering using the indep-pairwise command in Plink [64], with a window size of 500kb, a step size of 50kb, and an R^2^ threshold of 0.8. We used these LD-filtered SNPs to generate a PCA in Plink [64] as well as a genetic relatedness matrix (GRM) in GCTA [65]. Finally, to prepare for cis eQTL mapping, we extracted 1,950,183 LD-filtered SNPs that fell within 500kb of the transcription start or end site of protein coding genes.

### Testing for ancestry and treatment effects on gene expression levels

To identify genes for which gene expression was significantly predicted by ancestry within a given condition, we used linear models implemented in limma [62]. Specifically, for each of the 12 cellular environments, we ran the following model on the SV-corrected residuals for each gene:

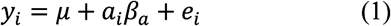

where *y*_*i*_ is the gene expression level estimate for sample *i, μ* is the intercept, *a*_*i*_ represents ancestry of the focal sample (AFR or EUR), *β*_*a*_ is the corresponding estimate of the ancestry effect, and *e*_*i*_ represents environmental noise.

To identify genes for which gene expression was significantly affected by a given treatment, we used a similar modeling approach again implemented in limma [62]. Specifically, for each of the 11 treatments, we ran the following model on the SV-corrected residuals for each gene:

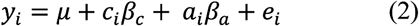

where all variables are as described above, with the addition of *c*_*i*_ which denotes the condition (treatment or control) and *β*_*c*_ which is the corresponding estimate of the treatment effect. After running both models 1 and 2, we extracted the p-value associated with the effect of interest (ancestry or treatment, respectively) and corrected for multiple hypothesis testing using a Storey-Tibshirani FDR approach [66]. A summary of the results of these analyses is provided in Tables S5 and S8. We also extracted the effect size and standard error estimates associated with the effects of interest for downstream analyses.

### Testing for cis eQTL effects on gene expression levels

We used the R package matrixeQTL [67] to test for cis eQTL (within 500kb) that affected gene expression variation in each cellular environment separately. Specifically, for each of the 12 environments, we ran the following model on the SV-corrected residuals for each gene:

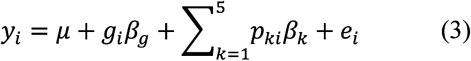

where *g*_*i*_ denotes the genotype of individual *i* in terms of number of copies of the minor allele (0, 1, or 2) and *β*_*g*_ is the corresponding estimate of the genotype effect. *p*_*ki*_ is the loading for principal component *k* for individual *i* (from a PCA on the filtered genotype matrix, as described above) and *β*_*k*_ is the estimate of the principal component effect. For each gene-SNP combination, we extracted the p-values, effect sizes, and standard error estimates associated with the genotype effect. We then used a Storey-Tibshirani FDR [66] to correct for multiple hypothesis testing. Results are summarized in Table S8.

We also ran a second analysis, using the same approach described above, but pooling the data across all 12 conditions. For this analysis, condition was also included as a covariate in the following model:

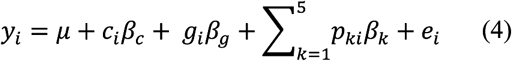

Finally, we note that we explored an alternative analysis strategy, in which we regressed out 1-20 principal components from the normalized (but not SV-corrected) gene expression matrix before fitting the models described in equation 3 (as in [9,30]). We did not find that this approach consistently increased our power to detect eQTL across the 12 cellular environments, and therefore opted for using the SV-corrected data in which a consistent pipeline could be applied to the full dataset.

### Estimating sharing of ancestry, treatment, and genotype effects

To understand the degree to which treatment, ancestry, or genotype effects mapped across different conditions have shared versus context-dependent effects, we used the empirical Bayes approach implemented in the R package mashR [20]. While previous studies have instead used linear models to 1) test for interaction effects between condition and genotype or ancestry or 2) test for ancestry or genotype effects on the fold change in gene expression levels estimated between treatment and control conditions [9,16,26], joint analysis via mashR is more appropriate to our design and provides many advantages. mashR is explicitly designed for quantitative assessment of effect size heterogeneity across conditions, increases power via joint analysis, and exploits patterns of similarity to provide improved estimates of effect size in each condition. It also provides a common framework for comparing effect sizes across many conditions, rather than relying on the comparison of many different “significant” lists derived from arbitrary p-value or FDR cutoffs. Similar meta-analytic approaches have been successfully applied to other high-dimensional datasets, such as GTEx [30]. We used similar pipelines to estimate sharing of treatment, ancestry, and genotype effects using mashR [20], with small modifications appropriate to each predictor variable.

First, using our multi-treatment (n=11) estimates of environmental effect sizes, we evaluated effect size concordance between all pairwise combinations of treatment-control pairs. Here, we followed the pipeline provided by the mashR authors for datasets that use the same reference or control condition samples across multiple comparisons (https://stephenslab.github.io/mashr/articles/intro_correlations.html). Specifically, we corrected for correlations among conditions in our data, and then used a combination of canonical and data-driven covariance matrices to fit the mashR model. From the mashR output, we extracted the posterior mean effect size and local false sign rate (LFSR) estimates. Following the authors’ recommendations, we considered a gene to have “shared” treatment effects across an arbitrary number of treatment if it has a local false sign rate (LFSR)<10% for at least one treatment and posterior effect sizes of similar magnitude (within a factor of two) for the other treatments. We always used the treatment with the lowest LFSR as the reference for the effect size comparison (we note that similar results were obtained when we used the median effect size across all treatments with LFSR<10%). In cases where the treatment effect was not shared across all 11 treatment-control pairs (aka “ubiquitous”), we considered the treatment effect to be “context-dependent”. Finally, we considered a gene to exhibit a special case of context-dependency, namely “condition-specific” effects, if it had evidence for treatment effects at a LFSR<10% and the posterior effect size estimate was not within a factor of two of any other treatment.

Second, using our multi-condition (n=12) estimates of ancestry effect sizes, we evaluated effect size concordance between all pairwise combinations of conditions. To do so, we followed the standard pipeline provided by the mashR authors (https://stephenslab.github.io/mashr/articles/intro_mash_dd.html) which uses a combination of canonical and data-driven covariance matrices to fit the mashR model. From the mashR output, we again extracted the posterior mean effect size and LFSR estimates, and used the same approach described above to identify ubiquitous, context-dependent, and condition-specific effects.

Finally, we assessed effect size sharing for cis eQTL mapped across all 12 cellular environments. In this case, there were too many tested gene-SNP pairs to evaluate all of them in the mashR framework (n= 8,109,941), so we followed the authors’ recommendations and focused on 66,614 gene-SNP pairs with some evidence for cis eQTL from matrixeQTL [67] (FDR<10% in at least one condition). We followed the pipeline suggested for eQTL (https://stephenslab.github.io/mashr/articles/eQTL_outline.html) and used a combination of canonical and data-driven covariance matrices derived from 50000 randomly chosen gene-SNP pairs to fit the mashR model. We then computed posterior summaries for the 66,614 gene-SNP pairs of interest using the model fit to randomly selected data. Finally, we extracted the posterior mean effect size and LFSR estimates, and used the same definitions described above to identify ubiquitous, context-dependent, and condition-specific treatment effects. In some cases, we also summarized our eQTL results at the gene rather than the SNP level; in these cases, we report the number of unique genes that have at least one eQTL that is shared between a given number of conditions (12=ubiquitous and 1-11=context-dependent). We note that with these definitions a given gene can be reported in more than one category.

A summary of the mashR treatment, ancestry, and cis eQTL analyses are provided in Tables S5 and S8. All estimates of LFSR and posterior mean effect sizes are available on Github.

### Testing the degree to which ancestry and genotype affect the response to treatment

We also used mashR to understand whether 1) individuals of African versus European ancestry responded differently to a given treatment and 2) genotype affects the response to a given treatment, as evidence for both phenomena has been shown in previous work [9]. To test point #1, we asked whether the posterior mean estimates of the ancestry effect were different between the treatment and control conditions for all 11 treatments; if so, this would indicate an effect of ancestry on the response to a given treatment. We used the same pipeline described for estimating sharing of treatment effects, and we considered a gene to exhibit differential responses to treatment as a function of ancestry if the LFSR was <10% in either the treatment or control condition (or both), and the posterior mean effect size estimates were not within a factor of two of one another. Using this approach, we found no evidence for ancestry effects on the response to treatment (except for 1 gene in the IGF-1 dataset). To validate this result, we also ran models in limma [62] that included an explicit interaction effect (*β*_*cxa*_) between treatment and ancestry. Specifically, for each of the 11 treatments, we ran the following model on the SV-corrected residuals for each gene:

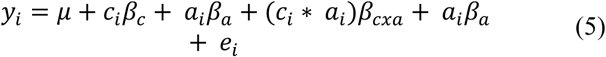

These results agreed with our results derived from mashR and revealed no significant ancestry x treatment effects at a 10% FDR. To test point #2, we asked whether the posterior mean estimates of the genotype effect were different between the treatment and control condition for all 11 treatments; if so, this would indicate an effect of genotype on the response to a given treatment. We used the same pipeline described for estimating sharing of cis eQTL effects, and we considered a gene to exhibit differential responses to treatment as a function of genotype if the LFSR was <10% in either the treatment or control condition (or both), and the posterior effect size estimates were not within a factor of two of one another. We found several thousand response eQTL for each treatment, and these results are summarized in Table S5.

### Enrichment analyses of treatment- and ancestry-associated genes

We used gene set enrichment analyses (GSEA) [68] to ask whether certain biological pathways were overrepresented among the set of genes that exhibited the strongest evidence for 1) differential expression in response to a given treatment and 2) ancestry-associated differences in expression. For #1, we sorted our gene list by effect size (output from limma) for each treatment effect separately and ran GSEA. For #2, we sorted our gene list by median effect size across all conditions, because very few genes exhibiting evidence for ancestry effect size heterogeneity across conditions. We assessed the significance of pathway enrichment scores via comparison to 1000 random permutations of gene labels across pathways, and controlled for multiple hypothesis testing using a Storey-Tibshirani FDR approach [66]. Results are reported in Figure S2, Tables S6, and Table S9.

We also tested whether ancestry-associated genes shared between ≥2/3 of all conditions (as determined by mashR [20]) were enriched within genes associated with 114 complex traits and diseases. To do so, we followed the approach of [8] and drew on publicly available results from Probabilistic Transcriptome Wide Association Studies (PTWAS) [21]. PTWAS combines eQTL data from GTEx [30] and GWAS data from several studies to identify genes that are likely along the causal pathway for a given complex trait or disease. We used hypergeometric tests to test for enrichment of ancestry-associated genes within each PTWAS trait-associated gene set and an FDR approach [66] to correct for multiple hypothesis testing. Results are reported in Table S10.

### Relationship between FST and ancestry-associated gene expression variation

We were interested in testing the hypothesis that genetic variation contributes to the observed differentiation in gene expression between African and European individuals. To do so, we followed the approach of [9] and asked whether genes with ancestry effects exhibit higher FST values between African and European populations relative to non-ancestry-associated genes. To do so, we first calculate FST for all 7,205,850 variants in our pre-LD filtered genotype dataset. We then generated gene-specific estimates by averaging FST values for variants within 5 kb (upstream or downstream) of the transcription start site of a given gene. Next, we used a Wilcoxon signed-rank test to ask whether FST values differed between genes with no evidence for an ancestry effect (LFSR>10% in all conditions) and 1) genes with any evidence for an ancestry effect in any condition or 2) genes with evidence for an ancestry effect in at least 2/3 of conditions. In a second approach, we also used linear models to test whether the number of cellular environments a gene exhibited significant ancestry effects in was predictive of the gene’s average FST value.

To further understand the contribution of genetic variation and selection to population differences in gene expression, we compared PST values for ancestry-associated genes to FST estimates between African and European individuals. PST is a phenotypic analog of FST (and a proxy for QST), and comparisons between the two measures can thus provide evolutionary insight [24,25]. Specifically, PST > FST is interpreted as evidence for diversifying selection, indicating different local optima for different populations. In contrast, PST < FST signifies uniform selection (also known as homogeneous, convergent, or stabilizing selection) and PST = FST suggests that phenotypic divergence between populations mimics neutral genetic divergence and is thus largely controlled by genetic drift. We used the R package Pstat [69] to calculate PST for ancestry-associated genes identified in each condition (FDR<10%) as well as a random sample of 500 genes. We compared these values to the genome-wide average of FST estimates between African and European individuals included in our dataset (for whom we also had whole genome sequencing data, n=454; see Figure 2 and Figure S3). To estimate 95% confidence intervals for the mean genome-wide FST estimate, we performed 1000 replicates of bootstrap resampling.

### Estimating the heritability of gene expression levels

We used GCTA and the GRM derived from the dataset of filtered genotypes to estimate the heritability of gene expression levels in each cellular environment. We followed the pipeline recommended by the authors (https://cnsgenomics.com/software/gcta/#GREMLanalysis) and estimated heritability for each of 10,156 genes in each of the 12 cellular environments. To understand whether heritability changed as a function of the cellular environment, we used a Wilcoxon signed-rank test to ask whether mean heritability differed between each treatment-control pair (see Figure 3 and Table S5). Because we observed upward biases in heritability estimates for conditions with the smallest sample sizes, we also 1) repeated the same analysis after subsampling each environment to n=100 individuals (performing 5 independent subsamples) and 2) used linear models to ask whether there was a consistent difference in per-gene heritability estimates between treatment and control conditions controlling for sample size.

### Enrichment analyses of ubiquitous and context-dependent eQTL and eGenes

We performed several enrichment analyses to understand the biology and putative phenotypic impacts of SNPs and genes that exhibited ubiquitous (shared across all 12 conditions) and context-dependent (condition-specific or shared across 2-11 conditions) eQTL.

First, to investigate the cellular mechanisms involved in generating ubiquitous and context-dependent eQTL, we downloaded ATAC-seq data generated for 20 LCLs from Yoruba 1000 Genomes individuals [32]. These data were preprocessed and provided as count matrices noting the number of reads mapped to a given region (n=2,533,845 windows) for a given individual. To identify strong and repeatable regions of open chromatin, we normalized the count matrix using the function voom in the R package limma [62]; we then retained regions for which the average normalized read counts were in the upper quartile of the entire dataset, and lifted over the region coordinates from hg19 to hg38 using the UCSC liftOver tool [70]. Finally, we used bedtools [71] to calculate the proportion of ubiquitous and context-dependent eQTL that overlapped with LCL ATAC-seq peaks, and we compared these proportions to background expectations derived from counting the proportion of all tested SNPs that overlapped with LCL ATAC-seq peaks. We performed these analyses using hypergeometric tests.

Second, we asked whether ubiquitous or context-dependent eQTL genes were enriched within the set of eGenes identified in unstimulated LCLs by GTEx [30] (i.e., genes with at least 1 eQTL identified at a 10% FDR). To do so, we used hypergeometric tests to compare our list of ubiquitous or context-dependent eGenes to GTEx eGenes, after first filtering for expressed genes that were common to both datasets. We also used hypergeometric tests to ask whether context-dependent eGenes that were not identified as eGenes in GTEx, but were identified as eGenes in our study, were enriched for genes that were also differentially expressed in our study (suggesting that cell perturbations “reveal” new eQTL). For this analysis, we used a combined list of all genes that were differentially expressed in any condition (FDR<10% from the limma output).

Third, we asked whether ubiquitous or context-dependent eGenes were enriched within sets of genes associated with 114 complex traits and diseases via PTWAS [21]. Here, we performed separate hypergeometric tests for each complex trait and each eGene list, and corrected for multiple hypothesis testing with a Storey-Tibshirani FDR approach [66].

Fourth, we downloaded the list of genes that are considered to be loss of function, mutation-intolerant genes, as curated by ExAC [34]. We then used Fisher’s exact tests and asked whether ubiquitous or context-dependent eQTL genes were enriched within the total set of mutation-intolerant genes.

Fifth, we downloaded the GWAS catalog [33] and filtered for SNPs with p<10^−8^. We then used Fisher’s exact tests to ask whether ubiquitous or context-dependent eQTL loci were enriched within the GWAS catalog.

### Evolutionary analysis of ubiquitous and context-dependent eQTL and eGenes

We performed two sets of analyses to address the roles of positive and negative selection in maintaining ubiquitous and context-dependent eQTL. First, we obtained two publicly available estimates of sequence conservation: phyloP scores [72] and phastCons scores [73]. The phyloP score measures the evolutionary conservation at each individual alignment site, with a positive sign indicating conservation and slower evolution than chance expectations, while a negative sign indicates relaxed constraint or positive selection and faster evolution than expected by chance. The phastCons score measures the probability that each nucleotide belongs to a conserved element, with a higherphastCons score representing greater sequence conservation. We obtained the per-site phyloP and phastCons scores from the 100-way vertebrate comparison available via the UCSC Genome Browser [70]. Following the methods of [74], we averaged the per-site measures across all exons in each protein coding gene to obtain per-gene phyloP and phastCons scores. Finally, we compared the mean per-gene conservation scores for ubiquitous eGenes and context-dependent eQTL genes to non eGenes using a Wilcoxon signed-rank test.

Second, we investigated a role for positive selection by obtaining per-site estimates of the integrated haplotype score (iHS), a commonly used measure of within-population recent positive selection [35,36]. We obtained genome-wide iHS estimates for each of the 10 populations included in our study from [36], and identified putative selection candidates as loci that fell in the >99^th^ percentile of |iHS| values in ≥2 populations (as in [9,16]). We then used Fisher’s exact tests to test for enrichment of iHS outliers in our ubiquitous and context-dependent eQTL sets, as well as the sets of response eQTL identified in each condition separately. Finally, we grouped the results of the response eQTL enrichment analyses into three treatment categories (immune stimuli, hormones, and environmental contaminants/novel cell stressors; see Table 1) and used a linear model to ask whether the enrichment effect sizes consistently differed between the immune stimuli treatments and the other two treatment categories.

### Data and code availability

FASTQ files for all mRNA-seq samples, as well as processed data matrices used for analyses, have been submitted to NCBI’s Gene Expression Omnibus (GEO). These files will be made public following publication. Code used to analyze the data and generate figures is available at: https://github.com/AmandaJLea/LCLs_gene_exp.

## Acknowledgments

We thank all members of the Ayroles lab for their feedback and support during the completion of this project. We also thank Andrea Graham and members of the Graham lab for thoughtful input and discussion. We thank Aaron Wolf, Christina Hansen, and Edward Schrom for assistance with cell culture protocols and facilities.

