## Supplemental Methods and Figures for "Diverse environmental perturbations reveal the evolution and context-dependency of genetic effects on gene expression levels"

3. Present address: Department of Biological Sciences, Vanderbilt University, Nashville, Tennessee, USA

The PDF file includes:

Supplementary Methods

Supplementary Figures S1 to S5

Supplementary References

**SUPPLEMENTARY METHODS**

**Cells and cell culture prior to experimental treatments**

Lymphoblastoid cell lines (LCLs) were obtained for 544 unrelated individuals included in the 1000 Genomes study [1]. All cells were ordered from Coriell Institute, and a complete list of the included lines is available in Table S1. Live cultures were shipped overnight to Princeton University in randomized batches of 25 (Table S1). Two batches of 25 were shipped on Monday and Tuesday of a given week and processed in parallel over the following two weeks; after the two-week period, another two batches of 25 arrived (and so on).

Upon arrival of a given batch of 25 samples, cells were incubated overnight in the flasks they were shipped in (unopened) at 37°C with 5% CO2. On day 2, cells were pelleted, counted using Trypan blue stain and a Countess Automated Cell Counter, and resuspended at a density of 500,000 cells per mL of cell culture media (RPMI + 10% fetal bovine serum + 1% antibiotic). Cells were then checked and split every 48 hours until a total of 14 million cells (12 million for our experiments and 2 million for cryopreservation) were obtained for a given line, or until 11 days had passed, whichever came first. At this time, we seeded 1 million cells in 2.5mL of media in each well of a 12 well plate, using one plate per individual. In cases where we had <12 million cells on the day of seeding, we plated however many wells we could. The 12 well plates were then placed overnight in an incubator and experimental treatments were performed on the morning of the following day (see below).

**Cell treatments**

On the morning of the experiment, a given 12 well plate was removed from the incubator and 5 ul of each treatment was added to a predesignated well. Table S2 shows the concentrations used for each of the experimental treatments. These concentrations were derived from the literature and modified in some cases based on pilot experiments. We note that our goal here was to learn about the general properties of genotype x environment interactions, rather than to create a cellular model for any individual treatment; thus, in some cases we chose concentrations that provoked a gene expression response in our pilot experiments rather than concentrations that were physiologically realistic. Because our treatments were dissolved in either water or ethanol, we also included these two molecules as controls, for a total of 12 conditions.

Following the addition of all 12 treatment and control molecules, plates were returned to the incubator for 4 hours, after which we spun down the plates for 5 min at 500 g at 4C. We then removed the cell culture media and washed the cells twice in cold 1x phosphate-buffered saline, spinning each wash for 10 min at 500 g at 4C. Finally, we lysed cells in 400 ul of a homemade lysis buffer that was made with the following recipe: 100 mM Tris-HCl at pH 7.5, 500 mM LiCl, 10 mM EDTA at pH 8, 1% LiDS, 5 mM dithiothreitol (DTT), and 10% beta mercaptoethanol. All lysed cells were then frozen at -80C.

**RNA extraction and library preparation**

12 well plates were thawed in batches of 8 or 12, depending on whether we were performing RNA extractions for all 12 conditions or just for 8 conditions (which we did for a subset of individuals). Depending on the circumstances, 12 conditions from 8 cell lines or 8 conditions from 12 cell lines were plated on a 96-well plate, with 3 wells randomly left empty as negative controls. To avoid introducing new batch effects, the samples that were plated together on a given 96-well plate were randomly chosen from a given batch of 50 samples that were received and processed during the same two-week period.

We used 200 ul of each sample to extract RNA using Zymo’s Quick-RNA 96 kit, following the manufacturer’s instructions. RNA extractions were repeated for a given plate if there was evidence of contamination in the negative control wells. mRNA-seq libraries were prepared using the published TM3’seq protocol [2] and a CyBio FeliX liquid handling robot (Analitik Jena). After the final PCR step, 5 ul of each library derived from a given cell line (n=8 or 12) were pooled, cleaned using SPRI beads, and quantified with a Qubit fluorimeter. The line level pools were then equimolarly combined into a plate level pool, which was visualized and quantified on an Agilent TapeStation. The total dataset (n=5223 libraries) was sequenced across four runs of the Illumina NovaSeq platform. Each sample was sequenced to a mean depth of 2.199 ± 2.731 (SD) million reads using 100bp single end sequencing.

**Low level RNA-seq data processing**

Each sample was trimmed for low quality bases and adapter contamination using cutadapt [3]. Trimmed reads were then mapped to the human reference genome (hg38) using STAR [4]. Next, we extracted reads that mapped uniquely and counted the number of reads that overlapped each gene using HTSeq [5] and the GENCODE v25 GTF (<https://www.gencodegenes.org/human/release_25.html>). If a sample had fewer than 250,000 reads mapped to protein coding genes, we excluded it from further analyses. This filtering step left us with 3886 samples and the metadata for these samples is provided in Table S3. Matrices of read counts per filtered sample were assembled and exported for further processing.

We translated our raw count data into transcripts per million (TPM), and filtered the dataset to exclude non-protein coding genes as well as genes that were lowly expressed in all conditions (median TPM<2 in all 12 conditions). This left us with 10157 genes for analysis. Next, we used the voom function in the R package limma [6] to normalize the count data. To remove variance attributed to batch and other technical effects, we conducted surrogate variable analysis [7] on the voom-normalized data (protecting the variance associated with treatment (12-way factor) and population (AFR/EUR)). We then fit linear models in limma [6] and regressed out three surrogate variables, the number recommended by the num.sv function. As expected, the three surrogate variables were highly correlated with the first 3 principal components of the normalized gene expression matrix (Pearson correlation R=-0.944, -0.929, and 0.819, all p<10^-16^). They were also correlated with known technical effects such as total read depth (R between SV1 and SV2 and total read depth=-0.319 and 0.131, both p<10^-16^) and sequencing batch (ANOVA of SV2 and sequencing batch: F=109.47, p<10^-16^). Importantly, because cell lines were randomized across all batch effects within the experiment, there should be no confounds between batch effects and our variables of interest. Principal components analyses of the SV-corrected dataset are presented in Figure S5.

**SUPPLEMENTARY FIGURES**

**Figure S1. Gene set enrichment analyses reveal changes in expected gene categories.** Results from gene set enrichment analyses testing for overrepresentation of particular gene ontology categories among differentially expressed genes in response to A) Tunicamycin, B) IFNγ, C) Gardiquimod, and D) B cell activating factor. We focus on these 4 environmental treatments as quality control, because we have relatively strong expectations for which biological pathways they should affect. In all cases, only the top 15 most significant categories are shown. Enrichment maps were created with the emapplot function in the R package enrichplot.

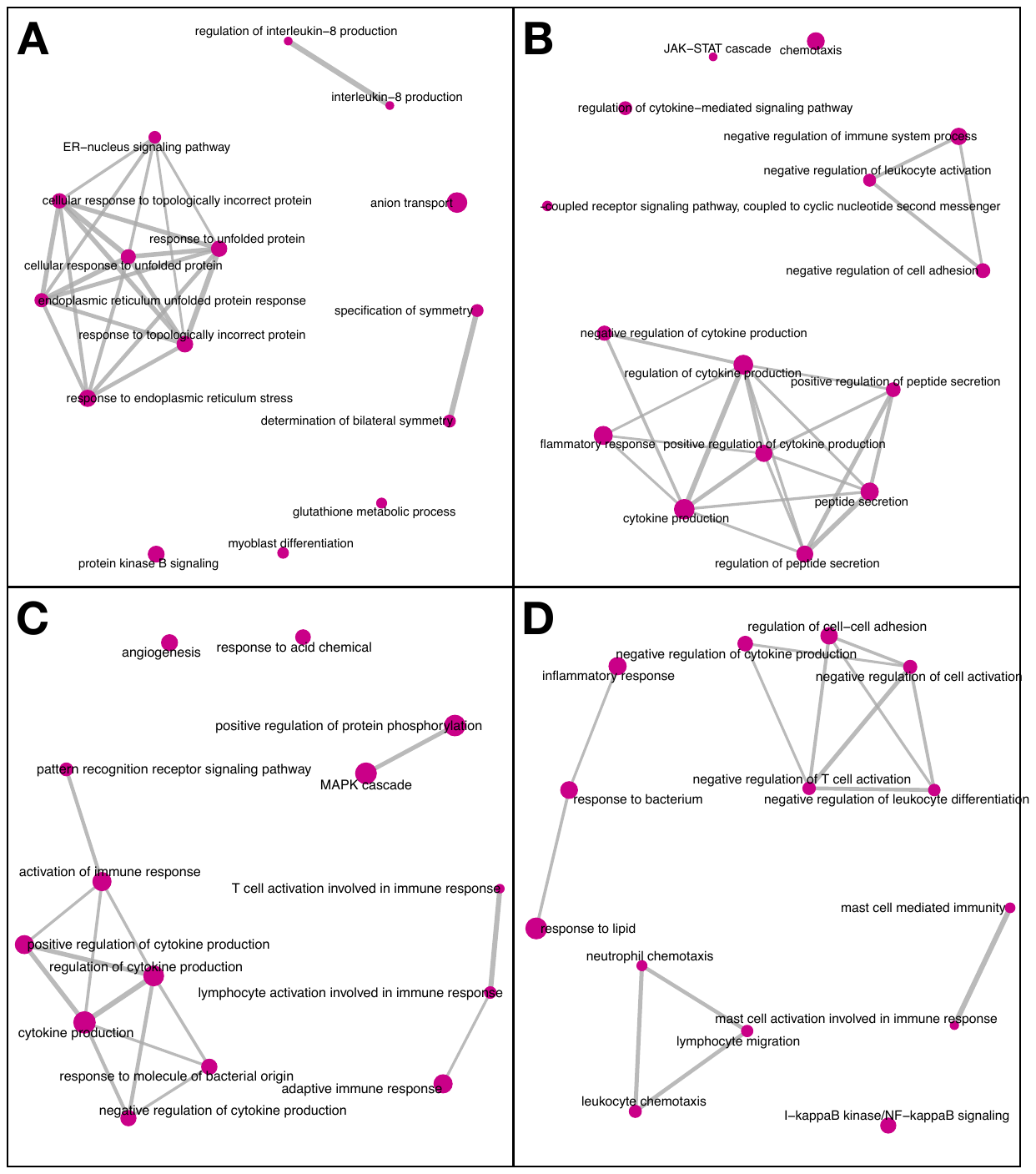

**Figure S2. Sharing of environmental and ancestry effects on gene expression levels.** Plots show the degree of correlation between the mashR [8] posterior mean estimates for (A) differentially expressed genes in response to a given environmental treatment and (B) differentially expressed genes as a function of ancestry (African versus European) in a a given cellular environment.

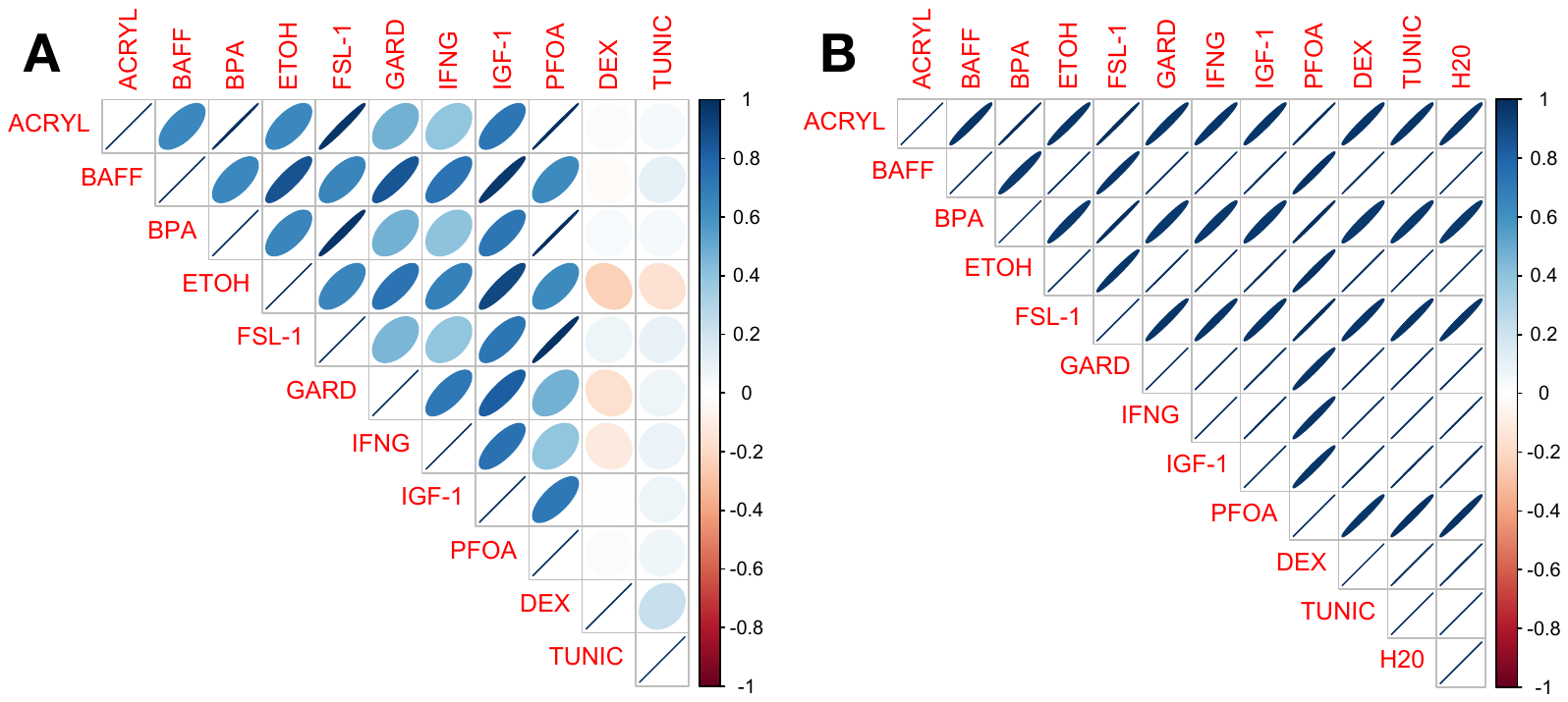

**Figure S3.** **Phenotypic differentiation in gene expression (P_ST_) versus genetic differentiation (F_ST_) for African versus European samples**. Plots show the distribution of P_ST_ values for: 1) AA genes identified in a given cellular environment (blue) and 2) a same-sized set of randomly selected genes (grey). The mean genome-wide F_ST_ value comparing genetic divergence between African and European samples is noted on the x-axis with an arrow. We find that all AA genes exhibit P_ST_ > F_ST_ in all cellular environments, indicative of diversifying selection [24,25].

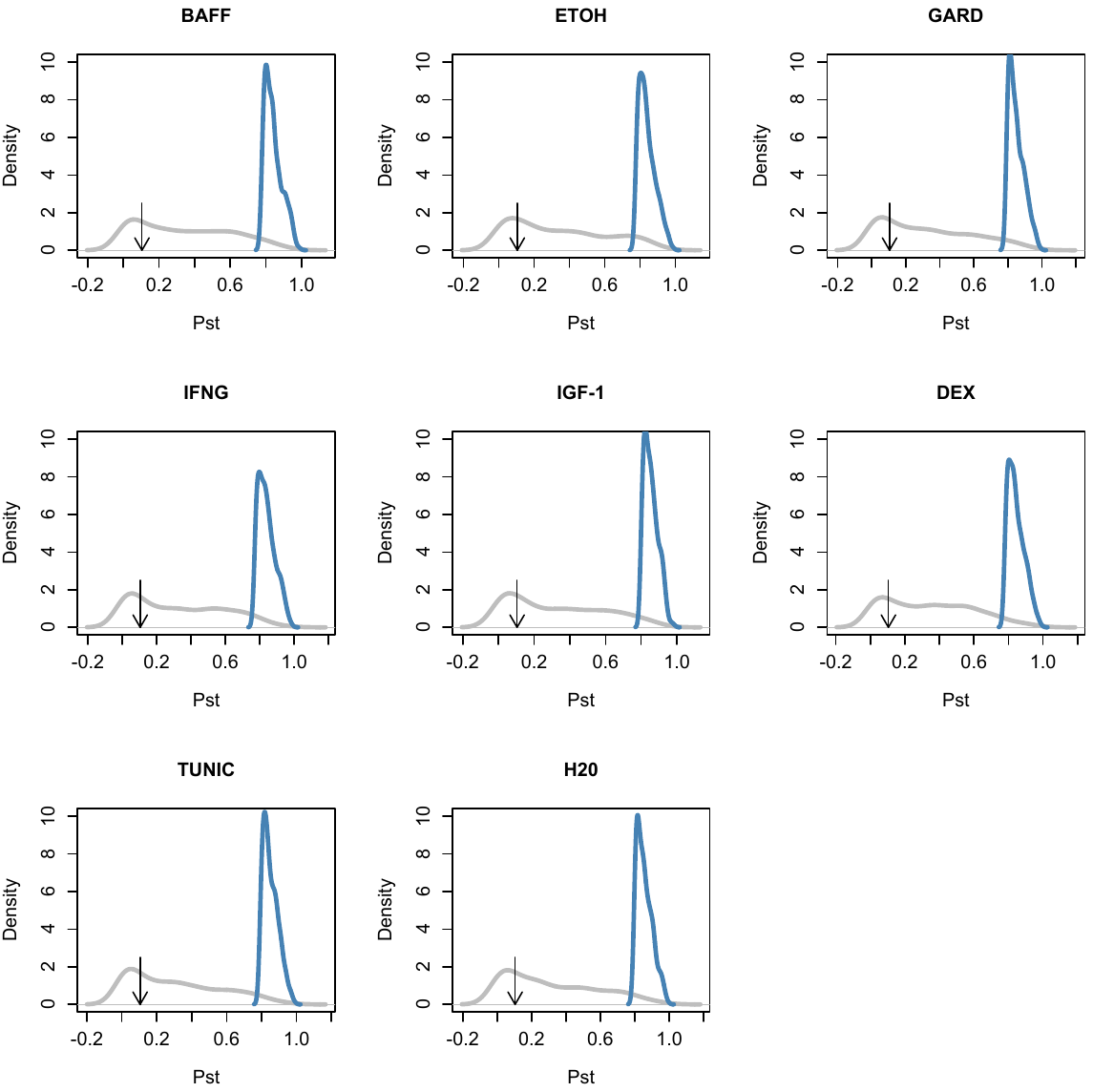

**Figure S4. Principal components analyses reveal treatment and ancestry effects.** We performed principal components analysis on the full, filtrered gene expression dataset (after surrogate variables were regressed out to remove technical effects). Top panels show the p-value from an analysis of variance testing for (A) treatment or (B) ancestry (African (AFR) versus European (EUR)) effects on the top 20 principal components (PCs). The dashed red line represents a nominal p-value of 0.05. Relationship between (C) treatment or (D) ancestry and principal component loadings for the PCs that were more strongly correlated with a given predictor variable.

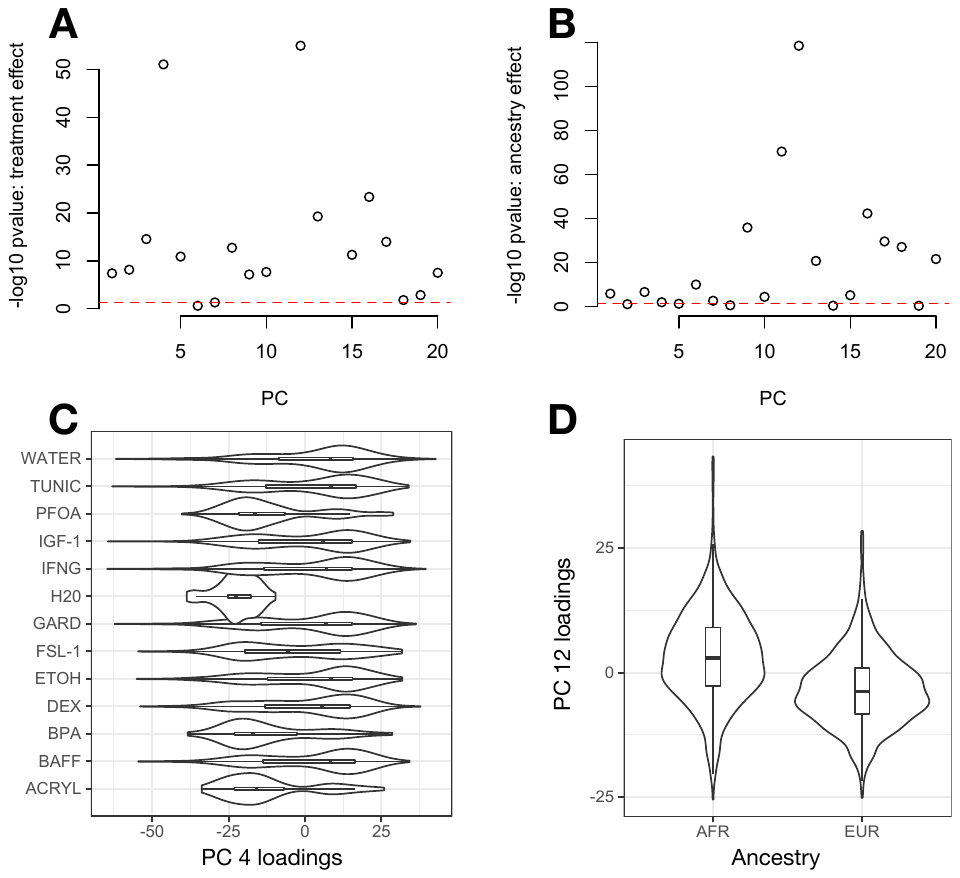

**Figure S5. Differentially expressed (DE) genes shared between environments.** (A) In Figure 1C, we present a version of panel A that shows that many genes are environment specific, few genes have similar effect sizes in 2 environments, and many genes have effect sizes that are similar across 3 or more environments. This result is driven by a large number of DEX-specific genes. Here we show the same distribution as plotted in Figure 1C, but with DEX excluded from the dataset. (B) For each environment, the number of genes included in the N=3 bar in Figure 1C is plotted. In combination with Figure 1D, this result emphasizes that many genes in our dataset have similar DE effect sizes shared across the group of 3 environmental contaminants (ACRYL, BPA, PFOA), or shared across some subset of a group of hormonal and immune treatments (GARD, IFNG, BAFF, IGF-1).

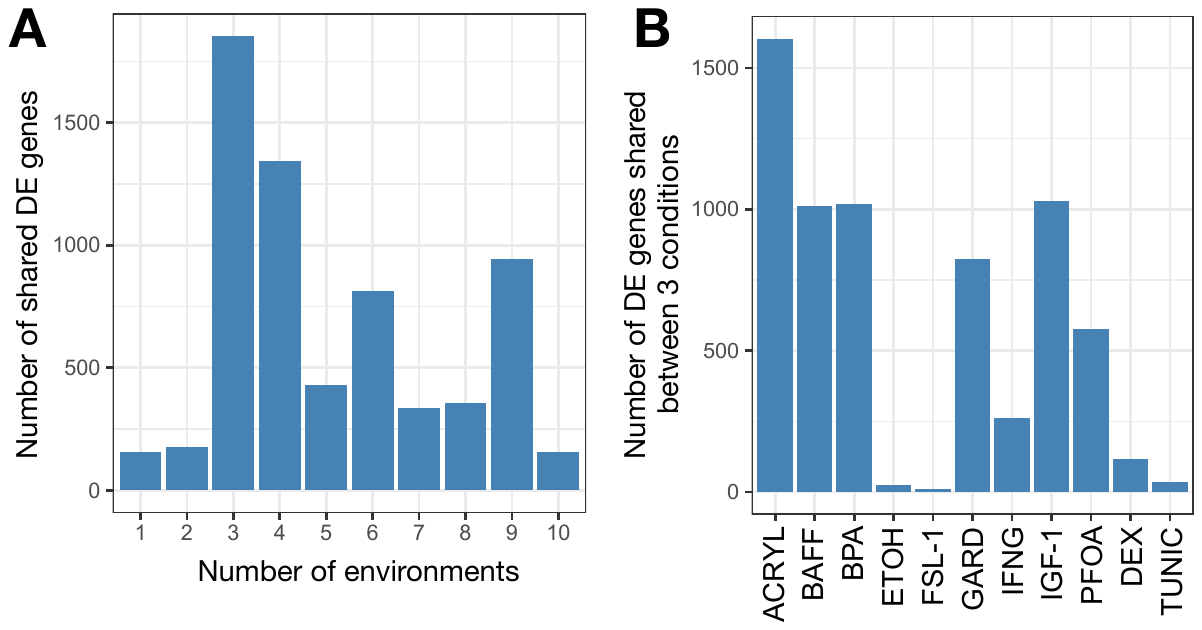
